# Bidirectional action of nitric oxide on mitochondrial respiration and permeability transition pore induced by calcium and palmitoylcarnitine

**DOI:** 10.1101/443986

**Authors:** Vladimir V. Dynnik, Elena V. Grishina, Nadezhda I Fedotcheva

**Affiliations:** Department of bioenergetics, Institute of Theoretical and Experimental Biophysics, Russian Academy of Sciences, Russia

**Keywords:** nitric oxide, mitochondrial respiration, permeability transition pore, mitochondrial nitric oxide synthase, guanylate cyclase and protein kinase G

## Abstract

The role of mitochondrial calcium-dependent NO synthase in the control of respiration and mitochondrial permeability transition pore (MPTP) opening, as well as possible involvement of mitochondrial NO synthase/guanylate cyclase/kinase G-signaling system (mtNOS-SS) in the regulation of these processes are not sufficiently studied. In this work, using rat liver mitochondria, we applied specific inhibitors of the enzymes of this signaling system to evaluate its role in the control of respiration and MPTP. The respiration was supported by pyruvate and glutamate or succinate in the presence of hexokinase, glucose and ADP. The results indicate that L-arginine and NO donors SNP and SNAP produce bidirectional concentration-dependent effects on the respiration and MPTP opening evoked by calcium ions or D,L-palmitoylcarnitine. Maximal activation of respiration was observed at 20 µM of L-arginine or SNP. At low concentrations, L-arginine (to 500 µM) and NO donors (to 50 µM) increased the threshold concentrations of calcium and D,L-palmitoylcarnitine required for the dissipation of the mitochondrial membrane potential and pore opening. The application of the inhibitors of NO synthase, guanylate cyclase, and kinase G eliminated both effects. These data indicate the involvement of mtNOS-SS in the activation of respiration and deceleration of MPTP opening. At high concentrations, L-arginine and NO donors inhibited the respiration and promoted pore opening, indicating that the inhibition induced by NO excess dominates over the protection caused by mtNOS-SS. These results demonstrate that the functioning of mtNOS-SS might provide a feedforward activation of respiration and a lowering of MPTP sensitivity to calcium and palmitoylcarnitine overload.

## Introduction

Numerous studies clearly demonstrate that exogenous nitric oxide (NO) suppresses mitochondrial respiration by inhibiting cytochrome c oxidase (COX) and complexes I and II of the electron transport chain [1–5]. The activation of calcium-dependent mitochondrial NO synthase (mtNOS) by its substrate L-arginine [6,7] or by Ca^2+^ plus L-arginine [8] also causes a sharp rise in the production of mitochondrial NO, which is followed by the inhibition of oxygen consumption [7].

In contrast to unidirectional action of NO on mitochondrial respiration, contradictory effects of NO on mitochondrial calcium retention capacity (CRC), mitochondrial permeability transition pore (MPTP) opening, and mitochondrial cytochrome c (CytC) release have been demonstrated over the last two decades. As early as 1999, it was shown that the activation of 2 mtNOS by Ca^2+^ and L-arginine induced CytC release, while the inhibition of mtNOS diminished it, raised mitochondrial potential (ΔΨm) and CRC [9]. Later it was demonstrated that NO evoked concentration-dependent effects on pore opening and CytC release [10]. It was shown that, being added at very low or high concentrations, the NO donor SpermineNONOate promoted mitochondrial swelling, CytC release, and MPTP opening induced by calcium, whereas in intermediate concentrations this compound caused protective effects. The protective and adverse effects of NO donors were attributed to possible action of S-nitrosothiols and peroxynitrite, correspondingly [9,10].

However, opposite to these results, ensuing studies indicated that the inhibitors of mtNOS promoted, while NO donors prevented the dissipation of ΔΨm and mitochondrial swelling induced by Ca^2+^ in isolated mitochondria [11]. Wherein, accumulated S-nitrosothiols were considered as final mediators providing the prevention of MPTP opening.

The concentration-dependent effects of NO donors were also demonstrated on permeabilized cells. It was shown that NO donors dose-dependently diminished mitochondrial Ca^2+^ uptake and, being applied in high doses, promoted MPTP opening [12,13]. It was assumed that the inhibition of Ca^2+^ uptake by intramitochondrial NO may represent negative feedback, which could prevent Ca^2+^ overload and MPTP opening [12]. According to another point of view, the inhibition of mitochondrial Ca^2+^ accumulation was explained by mitochondrial membrane depolarization and fall of ΔΨm induced by NO [14].

Modern experiments also demonstrate that moderate doses of nitroglycerine increase CRC and prevent Ca^2+^-dependent MPTP opening. NO and reactive nitrogen species are considered as the mediators, which may improve mitochondrial calcium handling and suppress pore opening [15].

Diverse effects of NO donors and L-arginine on mitochondrial respiration, ΔΨm, and MPTP are generally explained by the mechanisms based on the redox regulation of mitochondrial processes with the involvement of S-nitrosylation [16–18] and S-glutationylation [19–22] of numerous proteins.

However, potential intramitochondrial mechanisms of protection may also include some signaling chains involved in calcium and NO interplay. Recently Seya and coauthors discovered that cardiac mitochondrial protein fraction possesses PKG activity [23] and provides cGMP synthesis triggered by NO donors [24]. Besides, the hydrolysis of cGMP by mitochondrial cyclic nucleotide phosphodiesterase PDE2A was demonstrated in brain and liver mitochondria in independent experiments [25]. All these data indicate that the elements Ca^2+^-dependent mtNOS/ GC/PKG-signaling system (mtNOS-SS) may operate in mitochondria. However, according to the results presented by Seya and coauthors [23], SNAP or 8-Bromo-cGMP induced calcium-dependent CytC release and apoptosis, while the inhibitors of NOS, GC, and PKG prevented these effects. These results contradict to the generally admitted pro-survival action of cytosolic NOS/GC/PKG1-SS directed at the prevention of MPTP opening and cell death [26–29].

MPTP is considered as a common pathway leading to the development of apoptosis and necrosis, which are observed in myocardial ischaemia-reperfusion (I/R) [28–34], acute steatohepatits and oxidative stress [31,35,36], and under the action of various drugs and toxins [31,37]. Calcium and reactive oxygen species (ROS) are recognized as key mediators involved in MPTP opening [31,32]. However, I/R and some other pathologic processes are characterized not only by a steep rise of Ca^2+^ and ROS but also by the accumulation of long-chain fatty acids [38–40] and their carnitine derivatives [38], which are often considered as triggering agents evoking mitochondrial calcium overload and oxidative stress [31,40,41]. Long-chain fatty acids and acylcarnitines are oxidized in mitochondria as long-chain acyl CoA’s, which inhibit various enzymes including NAD(P)H-dependent dehydrogenases [42–44]. Our preliminary results indicate that D,L-palmitoylcarnitine (PC) excess induces CsA-dependent dissipation of ΔΨm, which may be prevented by SNAP [45].

In this work, we investigated the involvement of mitochondrial Ca^2+^ →mtNOS→NO→GC→cGMP→PKG-signaling system (mtNOS-SS) in the control of mitochondrial respiration and MPTP opening. The experiments carried out on isolated rat liver mitochondria demonstrated that L-arginine and NO donors SNP and SNAP at low concentrations stimulated mitochondrial respiration and suppressed the dissipation of ΔΨm and MPTP opening evoked by calcium or PC excess. The inhibitors of mtNOS, GC, and PKG abrogated both effects, indicating the involvement of mtNOS-SS in the activation of respiration and deceleration of MPTP opening. We assume that these results may provide mechanistic insight into the implication of mtNOS-SS in the regulation of mitochondrial functions.

## Results

### NO donors and L-arginine evoke the activation or inhibition of mitochondrial respiration depending on their concentrations

Coupled rat liver mitochondria held State 4 respiration rate of about 19 ± 1.4 ng-at O/min/mg prot. in the presence of L-glutamate and pyruvate as substrates. The addition of ADP (in the presence of glucose and hexokinase) increased the oxygen consumption rate to steady state values VO2ss = 54.5 ± 1.7 ng-at O/min/mg prot. Representative polarographic traces (O_2_ traces) are depicted on Fig 1. Black traces correspond to control experiments, while coloured traces characterize the influence of low (blue) and high (red) concentrations of the NO donor SNP (Fig. 1A) and L-arginine (Fig. 1B) on mitochondrial respiration. The preincubation of mitochondria with low concentrations of SNP (20 µM; Fig. 1A) or L-arginine (20 µM; Fig. 1B) resulted in an increase of VO2ss. At high concentrations, SNP (400 µM; Fig. 1A) and L-arginine (1000 µM; Fig. 1B) caused the inhibition of mitochondrial respiration.

**Fig. 1.**
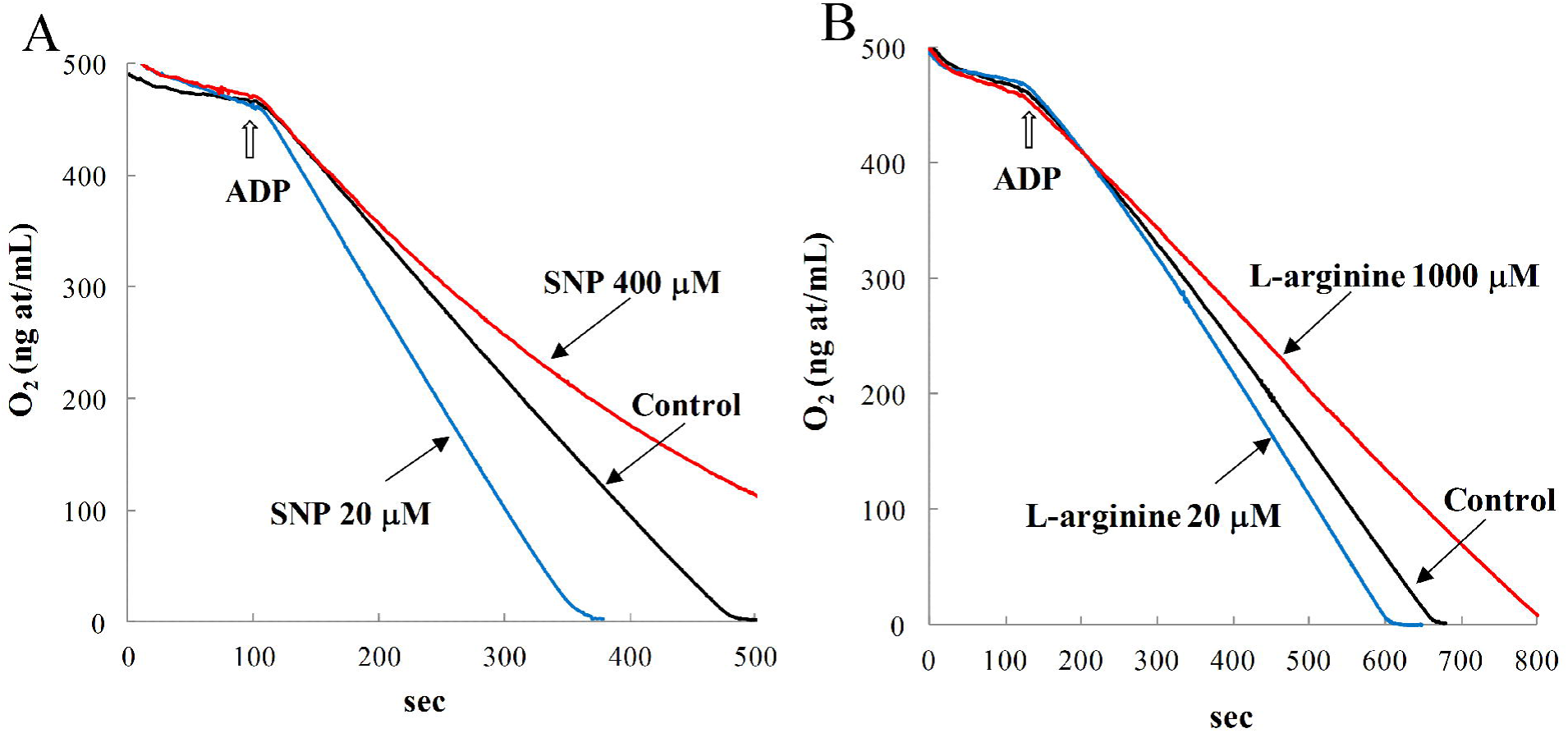
Activation and inhibition of mitochondrial respiration by NO donor SNP (left) and L-arginine (right). (A,B). Representative polarographic traces of mitochondrial respiration (O_2_ traces). Mitochondria (1.0 mg) were incubated in a closed chamber of 1 mL. The incubation medium included 0.5 mM pyruvate, 10 mM L-glutamate, 10 mM glucose and hexokinase. Steady state respiration was evoked by 0.75 mM ADP. Black traces describe control experiments. Blue traces correspond to low concentrations of SNP (A, 20 µM) and L-arginine (B, 20 µM) in the medium. Red traces match to high concentrations of SNP (A, 400 µM) and L-arginine (B, 1000 µM).

The calculated average values of VO_2_ss are depicted on Fig. 2. Left (black) columns correspond to control experiments. Two sets of violet and blue columns characterize the impact of different concentrations of SNP (Fig. 2A) and L-arginine (Fig. 2B) on mean values of VO_2_ss. The data show that the application of 10-50 µM SNP markedly activate steady state respiration rate. The maximal increase in VO2ss by 39% was observed at 20 µM SNP (Fig. 2A). In a like manner, VO2ss increased by 32% at low concentration of L-arginine (Fig. 2B, 20µM).

**Fig. 2.**
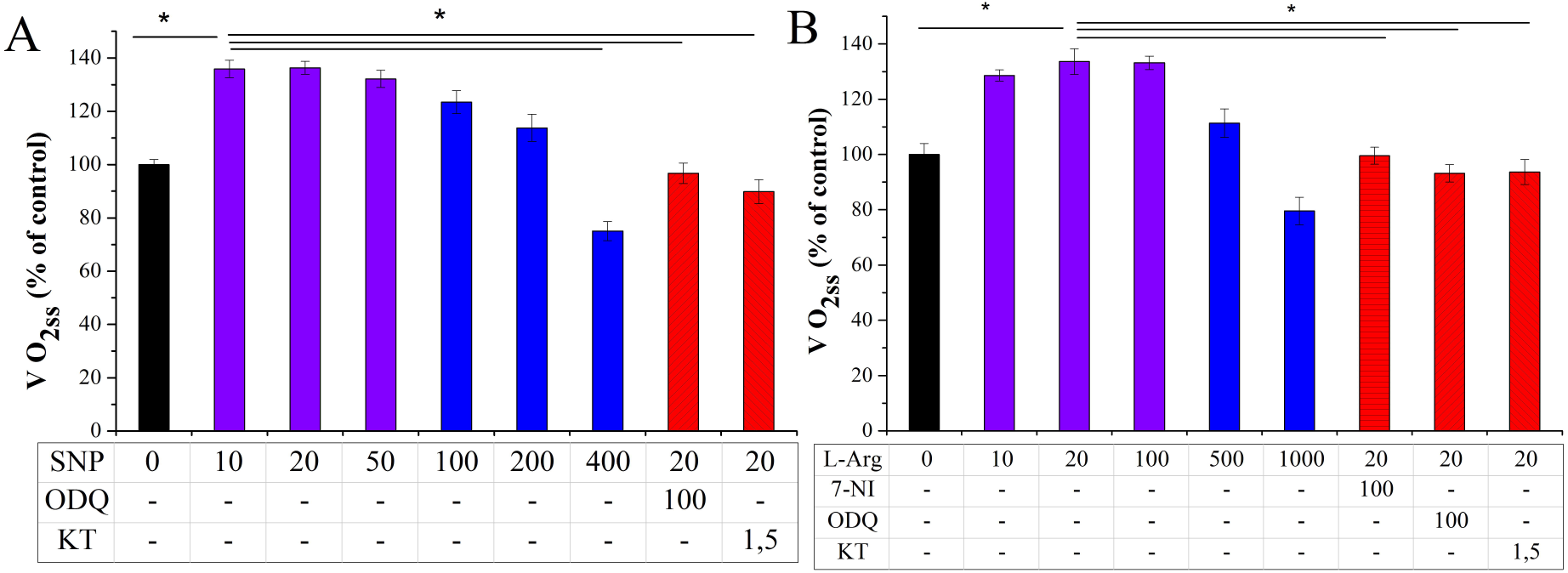
Bidirectional concentration-dependent effects of SNP (A) and L-arginine (B) on steady state respiration rate (VO_2_ss) and the influence of the inhibitors of mtNOS-SS on the values of VO2ss. The experimental conditions are the same as on Fig. 1. All columns represent the mean values of steady state respiration rates VO_2_ss. Black columns represent the data of control experiments (n = 4). Violet and blue columns describe the experiments performed with different concentrations of SNP (A) and L-arginine (B) in the medium (n = 5). Red hatched columns depict the impact of the inhibitors of NOS, GC, and PKG (7-NI, ODQ, and KT, respectively) on the values of VO_2_ss (n = 6). The values of VO_2_ss were calculated using the linear parts of corresponding polarographic traces at the concentrations of [O_2_] = 300-320 ng-at O/mL. The data are presented as the mean ± S.E.M. The concentrations of SNP, L-arginine, 7-NI, ODQ, and KT are given in µM. The control VO_2_ss value of 54.5 ± 1.7 ng-at O/min/mg protein was taken as 100%. Student’s *t*-test was applied. The compared pairs of VO_2_ss values are marked by horizontal lines placed above the columns. Symbol * indicate p < 0.05.

The activation of respiration by low concentrations of L-arginine or SNP was statistically significant in all experiments presented above (p < 0.05). Similar activation may be also evoked by low doses of NO donor SNAP, which stimulates VO_2_ss in the range of concentrations from 5 to 100 µM (not shown). Besides, the effects of NO donors and L-arginine do not depend on the substrates used and may be reproduced on mitochondria respiring with 1mM pyruvate and 5 mM L-malate or 5 mM succinate (in the presence of 2 µM rotenone) as substrates. In the last case, 10 µM SNP or 20 µM L-arginine raised VO_2_ss by 24% and 19%, correspondingly.

At high concentrations, SNP (400 µM) and L-arginine (1000 µM) diminished steady state respiration rate by 20–22% compared to control state (Figs. 2A, B). The inhibition of mitochondrial respiration by SNP (or L-arginine) represents a well known phenomenon [5–7], which may be based on competitive inhibition of COX by the excess of NO [1,6].

### Involvement of mtNOS-SS in the activation of mitochondrial respiration

Detected activation of the respiration by low concentrations of SNP, SNAP, and L-arginine contradicts the known inhibition of mitochondrial respiration by these substances [1–7]. We suggested that mtNOS-SS may be involved in the activation of mitochondrial respiration by SNP, SNAP, and L-arginine. In the next experiments we applied selective inhibitors of mtNOS-SS to evaluate the influence of this signaling system on the respiration. The results presented on Fig. 2 by red hatched columns demonstrate the effects of the inhibitors of NOS (7-NI), GC (ODQ), and PKG (KT). Figure 2A shows that ODQ and KT prevented the activating effect of 20 µM SNP by lowering VO_2_ss by 27% and 34%, respectively. Also, the activation of respiration by 20 µM L-arginine was not observed after the incubation of mitochondria with 7-NI, ODQ, and KT (Fig. 2B).

### Dissipation of the mitochondrial potential and inhibition of respiration by PC. Protection provided by low concentrations of L-arginine

Previously we have shown that high concentrations of PC (above 50 µM) induced a steep dissipation of ΔΨm and loss of mitochondrial calcium. In the presence of CsA (2 µM), critical concentration of PC (PC*) required for the dissipation of ΔΨm and pore opening raised several fold. Likewise, 50 µM SNAP increased PC* and substantially suppressed deleterious action of PC [45].

At low concentrations, added PC (to 20 µM) stimulated steady state rate of respiration VO2ss in mitochondria oxidizing pyruvate and L-glutamate by 30-35%. On the contrary, the application of 50 µM PC (PC*) triggered the dissipation of ΔΨm and the suppression of respiration (Fig. 3A, black traces). Preincubation of mitochondria with L-arginine (200 µM) prevented the dissipation of ΔΨm and recovered steady state respiration (green traces). The inhibitor of PKG KT abrogated protective effects of L-arginine by restoring the dissipation of ΔΨm and inhibition of the respiration caused by PC excess (red traces). This reveals the involvement of PKG, as final mediator of mtNOS-SS, in the protection provided by L-arginine.

**Fig. 3.**
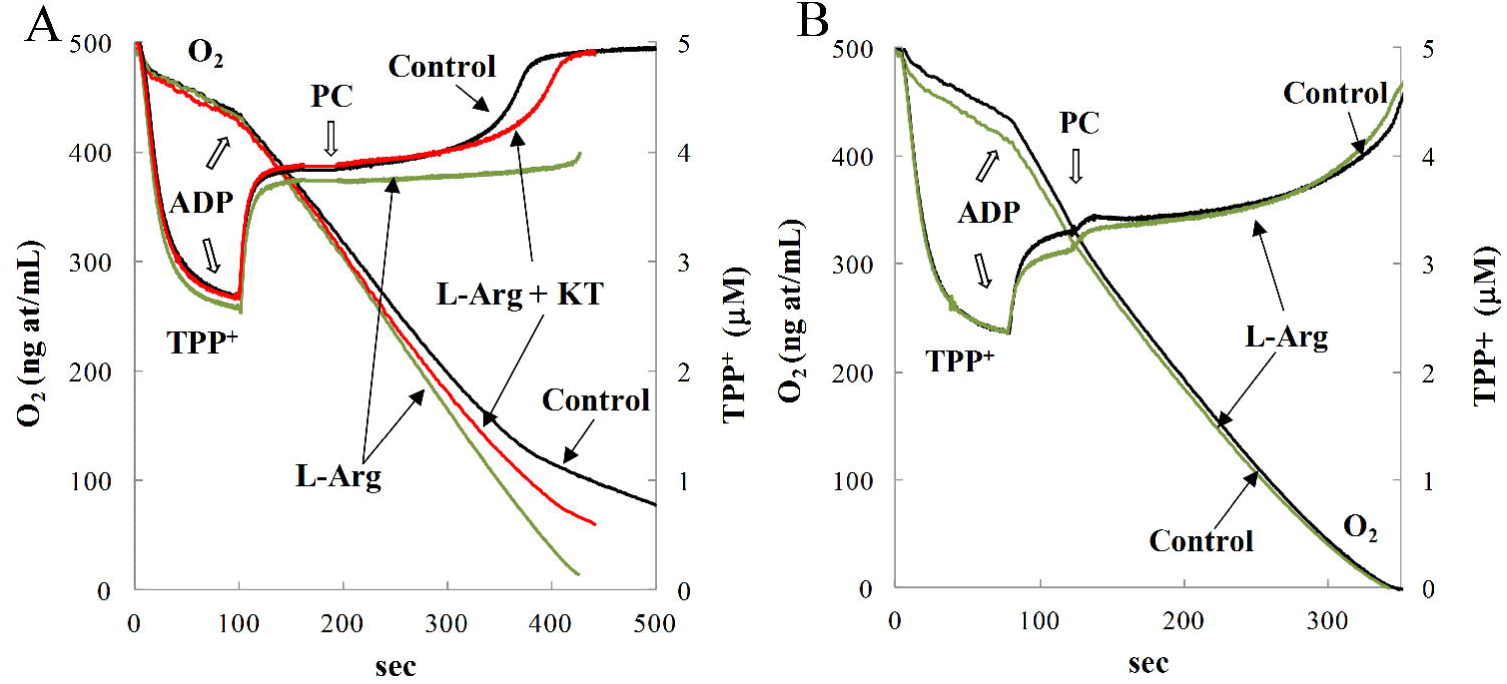
L-arginine prevents the dissipation of ΔΨm and suppression of respiration evoked by PC excess (A). The disappearance of the protective effect of L-arginine in mitochondria respiring on succinate plus rotenone (B). (A) Impact of PC excess (50 µM) and L-arginine (200 µM) on mitochondrial respiration (O_2_ traces) and ΔΨm (TPP^+^ traces). The experimental conditions are the same as on Fig. 1. PC was added 1 min after the activation of respiration by 0.75 mM ADP. Mitochondria were preincubated: without added L-arginine (control, black traces), with L-arginine (green traces), and with L-arginine plus 1.5 µM KT (red traces). (B) All conditions of the experiment correspond to those of panel A, except that incubation medium included 5 mM succinate as substrate plus 2 µM rotenone. L-Arg = L-arginine.

Importantly, L-arginine cannot prevent the dissipation of ΔΨm in mitochondria oxidizing succinate (Fig. 3B). Oxidation of succinate in the presence of rotenone provided high values of VO2ss (96 ± 2.8 ng-at O/min/mg prot.). However, in this case, all NAD(P)-dependent dehydrogenases and substrate level phosphorylation are not functioning and mitochondrial GTP may be synthesized only via transphosphorylation from ATP. Like to previous example (Fig. 3A), 50 µM PC induced the dissipation of ΔΨm and suppresed the respiration (Fig. 3B,black traces). But, L-arginine did not protect mitochondria against the deleterious effect of PC (Fig. 3B, green traces). The disappearance of the protective effect of L-arginine may be explained, assuming that at PC excess mitochondria oxidizing succinate cannot keep appropriate concentrations of NADPH and GTP required for functioning of mtNOS-SS.

### Impact of mtNOS-SS on the values of threshold concentrations of PC (PC*) required for the dissipation of ΔΨm

To evaluate critical threshold concentrations of PC* required for the dissipation of ΔΨm and MPTP opening, we used sequential additions of 20 µM PC (Fig. 4A). Control experiment is shown by brown line. In сontrol experiment third addition of PC (PC* = 60 µM) evoked a steep decrease of ΔΨm. SNAP evoked partial protection by raising the total concentration of PC to a new critical value of 100 µM.

**Fig. 4.**
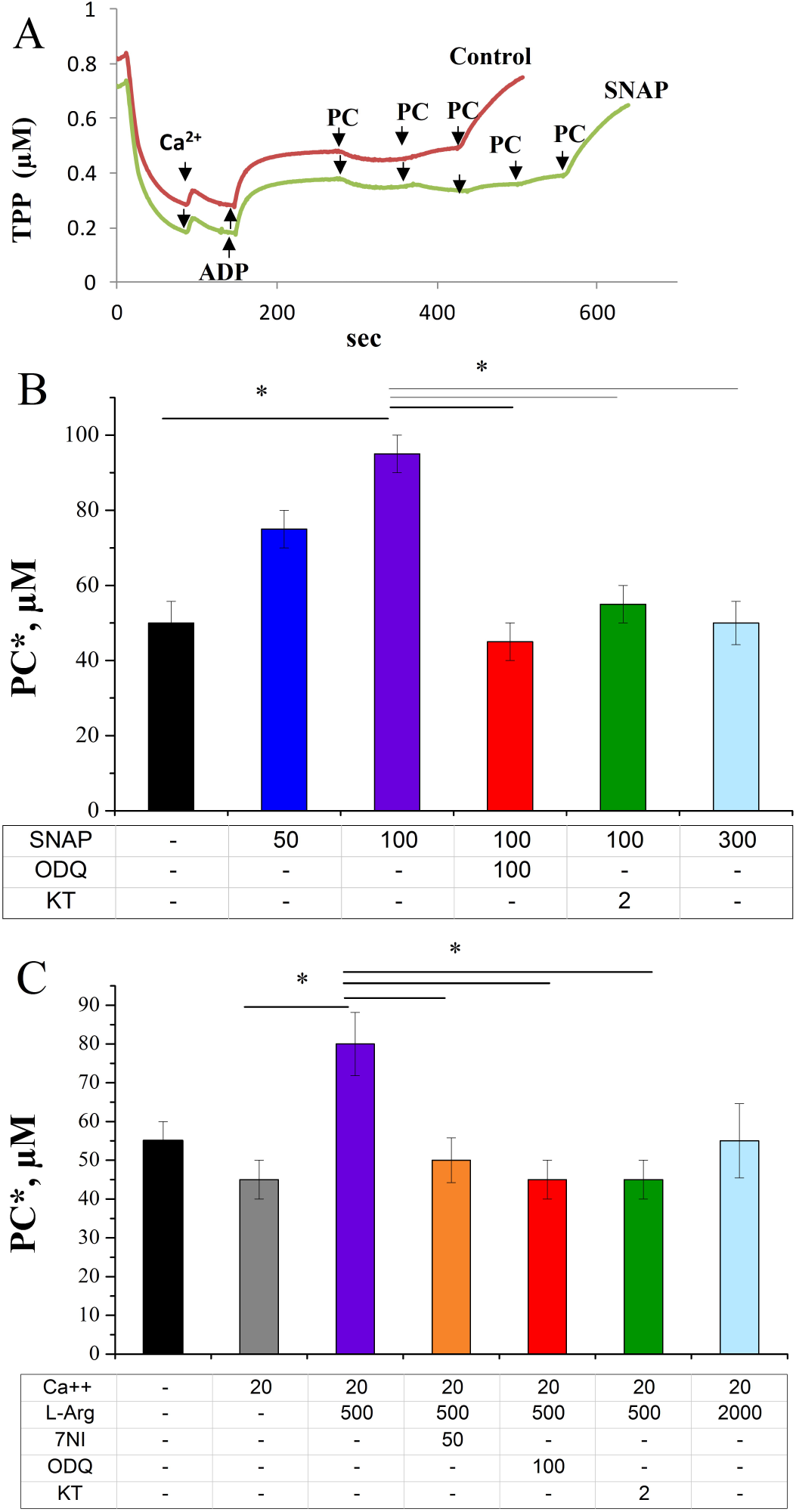
Impact of mtNOS-SS on threshold concentrations of PC* required for the dissipation of ΔΨm and MPTP opening. Mitochondria (1.0 mg of protein) were incubated in an open chamber of a volume 1 mL without SNAP (control, brown trace) or with 100 µM SNAP (green trace). The incubation media correspond to those on Fig. 1. Steady state respiration was set by 750 µM ADP. Ca^2+^ (20 µM) was added before the application of ADP. (A) An example of the determination of threshold PC* concentrations. The third drop of 20 µM PC evoked a fall of ΔΨm, which corresponds to the threshold value of PC* = 60 µM (Control, red line). Preincubation of mitochondria with SNAP raised PC* to 100 µM (green line); (B) The mean threshold values of PC* measured in control experiments (black columns) and in the experiments performed on mitochondria preincubated with SNAP (blue and violet columns), SNAP + ODQ and SNAP + KT (red and green columns); (C) The influence of 500 µM L-arginine on PC* value (violet) and elimination of its effect by the inhibitors of NOS, GC and PKG (7-NI, ODQ and KT; brown, red and green columns, respectively). All concentrations are given in µM. L-arg = L-arginine.Data represent mean ± S.E.M. n = 4. The compared pairs of PC* values are marked by horizontal lines placed above columns. Symbol * indicate p < 0.05.

Two diagrams presented on Figs. 4B,C indicate the involvement of mtNOS-SS in the regulation of MPTP sensitivity to PC. As shown on Fig. 4B, SNAP raised critical PC* level in a concentration-dependent manner, being most effective at the concentration of 100 µM. Preincubation of mitochondria with 100 µM SNAP lowered the sensitivity of MPTP to PC about two fold by increasing PC* level from control value of 50.0 ± 5.8 µM to 95.0 ± 6.2 µM (violet vs. black columns). The application of higher concentrations of SNAP (above 300 µM) was ineffective. The protection provided by low concentrations of SNAP was eliminated by the application of GC and PKG inhibitors ODQ and KT (Fig. 4B). As a result, the sensitivity of ΔΨm (and MPTP) to PC excess increased about two times and was close to control value of PC*.

Like SNAP, L-arginine diminished MPTP sensitivity to PC by increasing its critical PC* value (Fig. 4C). To strengthen the effect of L-arginine on the dissipation of ΔΨm by PC we included in the incubation medium the activator of mtNOS Ca^2+^. Calcium and PC act synergistically by reinforcing the effects of each other [45]. In the absence of Ca^2+^in the medium, the PC* value was equal to 55.0 ± 5.0 µM (black). The addition of 20 µM Ca^2+^ diminished PC* to 45.0 ± 5.0 µM (grey). At higher concentrations, Ca^2+^ evoked marked lowering of PC* (not shown). So, we used this boundary concentration of Ca^2+^ (20 µM) to attain substantial activation of mtNOS (of mtNOS-SS) by L-arginine. L-arginine (up to 500 µM) increased critical PC* level about two times compared to control value (violet vs. grey columns).This effect was eliminated by the inhibitors of NOS, GC and PKG, which returned critical PC* values to the range of control values.

Thus, the data presented at Fig. 4 demonstrate that the activation of mtNOS-SS by L-arginine or by NO switch on some new protective mechanisms directed on the suppression of ΔΨm dissipation and MPTP opening by PC excess.

### Impact of mtNOS-SS on mitochondrial calcium retention capacity

Calcium retention capacity (CRC) was used as second parameter, which may characterize the involvement of mtNOS-SS in MPTP control. The values of CRC were measured by the standard procedure of sequential loading the medium with 20 µM Ca^2+^ (CaCl_2_). Like to the experiments presented above (Fig. 4), PC (20 µM) was added to reinforce the effect of calcium on MPTP. The inhibitor of PKG KT (2 µM) diminished CRC in comparison with control value by 30 % (Fig. 5A). The diagrams presented on Figs. 5B,C demonstrate the involvement of mtNOS-SS in the control of CRC. Addition of 20 µM PC in the incubation medium evoked CRC lowering by19-20% (grey vs. black columns). This effect indicates calcium and PC interplay in the regulation of ΔΨm dissipation and pore opening and corresponds to their synergistic action presented on Fig. 4C.

**Fig. 5.**
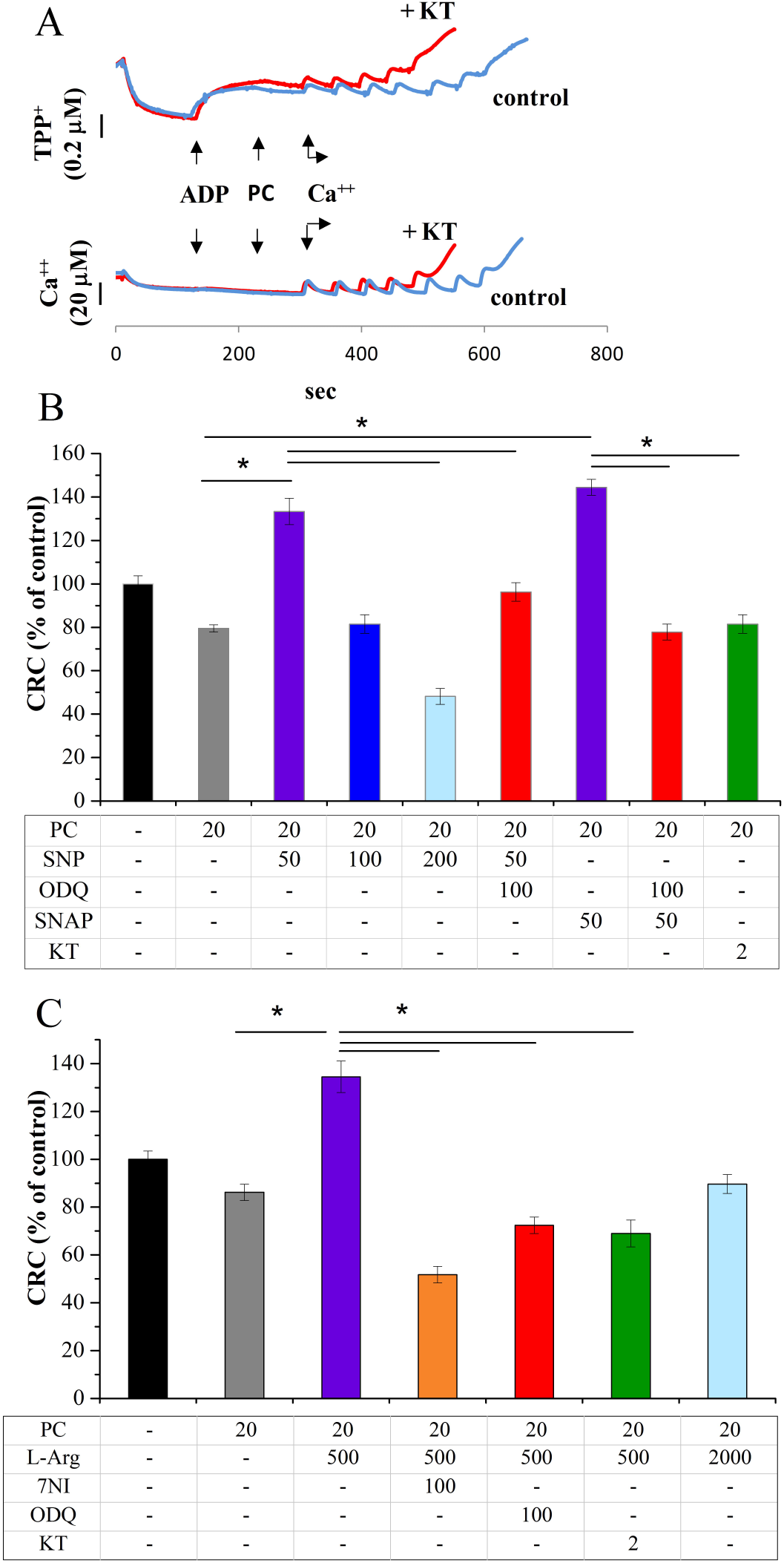
Involvement of mtNOS-SS in the regulation of mitochondrial CRC. (A) Determination of CRC by registering steep alterations of ΔΨm (TPP^+^, top traces) and Ca^2+^ concentration in medium (bottom traces). Mitochondria (1.0 mg) were incubated in an open chamber of 1 mL volume without PKG inhibitor KT (control, blue) or with 2 µM KT (red). The incubation media correspond to those on Fig. 1. PC (20 µM) was added after stimulation of respiration by 750 µM ADP. The seventh addition of 20 µM Ca^2+^ evoked an extrusion of Ca^2+^ from the matrix, dissipation of ΔΨm, and pore opening in mitochondria, which corresponds to CRC = 140 nmol/mg. The incubation of mitochondria with KT diminished CRC to 100 nmol/mg; (B) Protective effects of SNP and SNAP on pore opening induced by Ca^2+^ overload. Here are presented the mean values of CRC. Initial experiments (black column) were carried out with pyruvate and L-glutamate as substrates. PC (20 µM) was added before loading mitochondria with Ca^2+^ in all other experiments. The control value of CRC (calculated for grey column) was equal to 135.0 ± 5.0 nmol/mg and was taken as 100%. Colored columns correspond to the experiments, in which mitochondria were preincubated with SNP, SNAP and the inhibitors of GC (ODQ) and PKG (KT). (C): Positive impact of L-arginine on CRC values and elimination of this effect by 7-NI, ODQ and KT. All other conditions are as on Fig.5B. All concentrations are given in µM. The data represent the mean ± S.E.M. n=4. Symbol *indicates p < 0.05. Compared pairs of CRC values are marked by horizontal lines placed above the columns.

At low concentrations, SNP and SNAP (up to 50 µM) caused a protective effect by raising mean CRC values by 75% and 80%, respectively (Fig. 5B: violet vs. gray columns). Higher concentrations of SNP (100 and 200 µM) evoked opposite effect by diminishing CRC values two to three times, correspondingly.

The protection provided by SNP and SNAP (both 20 µM) was eliminated by the inhibitors of the enzymes of mtNOS-SS (Fig. 5B). The inhibitor of GC ODQ resulted in a significant decrease of CRC values for SNP and SNAP nearly to control values. The inhibitor of PKG KT produced a similar effect.

Like to the action of NO donors, 500 µM L-arginine raised the CRC value by 66% (Fig. 5C). The inhibitor of mtNOS 7-NI evoked a substantial decrease in CRC, which was about three times lower than maximal value of CRC. The inhibitors of GC and PKG also eliminated the protection provided by L-arginine. At high concentrations, L-arginine (2000 µM), like to NO donors excess (Fig. 5B, blue column), promoted pore opening by diminishing CRC below control values (Fig. 5C, blue column).

Thus, the data presented on Figs. 4, 5 demonstrate that at low concentrations L-arginine and NO-donors produce protective effects, which are characterized by marked rise of PC^*^ and CRC values in comparison with control values. Substantial decrease of both parameter values evoked by the inhibitors of mtNOS-SS indicates the involvement of this signaling system in the control of MPTP.

## Discussion

When analyzing the known contradictory effects of NO on the induction of MPTP by Ca^2+^ [9–15], we assumed that calcium-dependent mtNOS-SS may represent a missing link involved in mitochondrial Ca^2+^ and NO interplay and MPTP control.

The results presented above show that SNP, SNAP and L-arginine produce bidirectional concentration-dependent effects on mitochondrial respiration and MPTP opening triggered by calcium and PC excess. At low concentrations, NO donors (to 50 µM) and L-arginine (to 100 µM) caused a substantial activation of respiration by increasing VO2ss by 30–40 % (Figs. 2A, B). Besides, NO donors and L-arginine suppressed the dissipation of ΔΨm and MPTP opening by increasing the values of two regulatory parameters (CRC and PC*) characterizing MPTP sensitivity to calcium and PC overload.

The application of the inhibitors of NOS, GC and PKG eliminated both effects, indicating the involvement of mtNOS-SS in the activation of respiration (Fig. 2) and deceleration of MPTP opening (Figs. 4, 5). Mitochondrial PKG may act as a final mediator involved in the activation of respiration and MPTP control. It is known that mitochondrial protein kinase A activates the respiration by phosphorylating the enzymes of respiratory chain [46–49]. A similar mechanism of action can also be inherent in PKG.

Calcium-dependent activation of mitochondrial respiration by mtNOS-SS creates positive feedforward loop, which reinforces the regulation of oxidative phosphorylation by calcium ions and may balance direct positive effects of calcium ions on key dehydrogenases of the Krebs cycle [8]. Besides, the transmission of calcium signal to the final mediator mtPKG creates a negative feedforward loop, which might prevent MPTP opening induced by Ca^2+^ and PC overload.

At high concentrations, L-arginine (above 500 µM) and NO donors (above 100 µM) inhibited the respiration (Figs. 1, 2) and promoted MPTP opening (Figs. 3, 5). Thus, we may conclude that NO and mtNOS-SS have an opposite effects of on the respiration and MPTP control. Apparently, inhibitory impact of mitochondrial NO excess may overcome positive effects provided by mtNOS-SS.

It is worth to note that mtNOS-SS cannot be considered as a redundant element in the multi-level control of oxidative phosphorylation and MPTP. Being calcium-dependent, this signaling system may be involved in the functioning of several feedback and feedforward regulatory loops, two of which are directed toward the activation of mitochondrial respiration and MPTP control. In this case, mitochondrial PKG, along with cytosolic PKG1, may be considered as a final mediator of protection involved in MPTP control.

It is well known that a number of kinases and phosphatases are transported into mitochondria and have here, as the targets, multiple proteins, including the components of MPTP complex [27,49–52]. Special mechanisms provide the transport of these proteins [50]. We might suppose that GC and PKG are imported into mitochondria using like mechanisms. However, direct experimental data confirming the localization of PKG and GC inside of mitochondria are currently lacking. This issue requires further investigations.

## Materials and methods

All animal procedures were fulfilled in accordance with the EU directive 86/609/EEC and approved by the Ethics Committee at the Institute of Theoretical and Experimental Biophysics, RAS, Russia. Male (six-to eight-week-old) Wistar rats were kept under the same conditions in air-conditioned and ventilated rooms at 20–22°C with a 12 h/12 h light-dark cycle. All experiments were performed at 26°C. Liver mitochondria were isolated using standard techniques of differential centrifugation in the medium containing 300 mM sucrose, 1 mM EGTA, and 10 mM Tris-HCl (pH 7.4). Mitochondrial preparations were washed twice with the release medium containing no EGTA, resuspended in the medium of the same composition, and stored on ice as described earlier [45]. Mitochondria incubation medium contained: 125 mM KCl, 3 mM KH_2_PO_4_, 10 mM HEPES (pH 7.4), 0.5 mM MgCl_2_. The content of mitochondrial proteins was determined by the Lowry method with bovine serum albumin as a standard. Oxygen consumption in a mitochondrial suspension was determined by the polarographic method with a Clark type O_2_ electrode in a closed chamber of 1 mL containing 1.0-1.1 mg mitochondrial protein, under continuous stirring. The difference of electric potential on the inner mitochondrial membrane (ΔΨm) was measured by determining the redistribution of lipophilic cation tetraphenylphosphonium (TPP^+^) between incubation medium and mitochondria. The concentration of TPP^+^ [TPP^+^] in the mitochondrial incubation medium was recorded by a TPP^+^ selective electrode. Changes in calcium ion concentration in the incubation medium were recorded by a Ca^2+^ selective electrode (Nico, Moscow, Russia). Simultaneous registration of ΔΨm and Ca^2+^ was carried out in an open chamber of volume 1 ml containing 1.0-1.2 mg mitochondrial protein under continuous stirring.

L*-*glutamate (10 mM) and pyruvate (0.5-1 mM) were used as substrates to keep the turnover of the Krebs cycle, provide sufficient NADPH production, and preserve substrate-level phosphorylation and GTP synthesis in the Krebs cycle. In some experiments, mitochondrial respiration was supported by succinate (5 mM) or pyruvate (1 mM) and L-malate (5 mM). All experiments were performed in the presence of 10 U/ml hexokinase, 10 mM glucose, and 0.5 mM MgCl. Subsequent addition in the incubation media of 0,75 mM ADP provided a high steady state respiration rate (VO_2_ss), which was close to State 3 respiration rate (80–90% of VO_2_max). All values of VO_2_ss = d[O_2_]/dt were calculated on the linear parts of polarographic tracks at appropriate values of pO_2_ (as indicated in the corresponding legends).

The opening of the MPTP was registered as a loss of calcium buffering capacity (a steep rise in calcium in the incubation medium) and/or dissipation of mitochondrial ΔΨm. MPTP opening was induced by the sequential loading of the incubation medium with 20 µM Ca^2+^ (CaCl_2_) or D,L-palmitoylcarnitine (PC). In this way, the values of mitochondrial CRC and the threshold concentration of PC (PC*) were determined as total concentrations of added Ca^2+^ and PC required for pore opening. To investigate the impact of mtNOS-SS on MPTP opening, we selected these values of CRC and PC* as regulatory parameters characterizing the sensitivity of MPTP to Ca^2+^ and PC overload. Besides, to evaluate the involvement of mtNOS-SS in the regulation of mitochondrial respiration, we determined the rates of steady-state respirationVO2ss.

The activation of mtNOS-SS was produced by the application of L-arginine, NO donors, and Ca^2+^; 7-NI, ODQ, and KT5823 were used to inhibit mtNOS, GC and PKG, correspondingly.

All reagents were purchased from Tocris (UK) and Sigma (USA). S-nitrosoacetylpenacillamine (SNAP), 7-nitroindasole (7-NI) and 1H-[1,2,4]oxadiazolo[4,3-a]quinoxalin-1-one (ODQ) were dissolved in ethanol and KT5823 (KT)-in dimethyl sulfoxide (DMSO). The final concentration of DMSO was 0.1–0.2%. Sodium nitroprusside (SNP) was dissolved in water.

Statistical analysis of the experimental data was carried out by applying SigmaPlot 11. Student’s *t*-test. p < 0.05 was taken as the level of significance. The data are presented by columns as mean ± S.E.M of four to six independent experiments. Compared pairs of values are marked by horizontal lines placed above the columns. Symbol * placed above the lines indicate that p < 0.05.

## Supporting information

Graphical Abstract

## Author Contributions

VVD coordinated the project and wrote the manuscript with inputs from all authors. EVG and NIF performed the experiments, analyzed and discussed the data. NIF participated in the revision of the paper. All authors reviewed the results and approved the final version of the manuscript. Resources: EVG, VVD.

## Acknowledgments

We acknowledge Mary Simonova, Milija Galimova and Alex Sergeev for technical support.

This work was supported by grants from Russian Federation for basic research (RFBR): project N№ 14-04-01695 to VVD; project N14-04-01597 to EVG.

