## Supplementary material for "Bidirectional action of nitric oxide on mitochondrial respiration and permeability transition pore induced by calcium and palmitoylcarnitine": Graphical Abstract

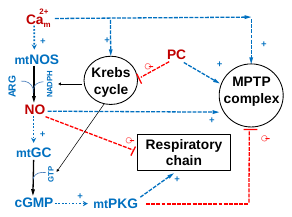

At low concentrations, L-arginine and NO donors activate respiration (+) and increase the threshold concentrations of Ca^2+^ and palmitoylcarnitine (PC) required for pore opening (+) through the activation of mtPKG. At high concentrations, NO overcomes the protection afforded by mtPKG, reinforces the inhibition of respiration (-), and promotes pore opening (-).
